# Alpha-to-beta- and gamma-band activity reflect predictive coding in affective visual processing

**DOI:** 10.1101/2021.06.03.446893

**Authors:** Andreas Strube, Michael Rose, Sepideh Fazeli, Christian Büchel

**Affiliations:** Department of Systems Neuroscience, University Medical Center Hamburg-Eppendorf, 20246 Hamburg, Germany

**Author notes:** Corresponding Author: Christian Büchel, Department of Systems Neuroscience University Medical Center Hamburg-Eppendorf 20246 Hamburg, Germany.

**Keywords:** Predictive Coding, Prediction Error, Expectation, Emotion, Arousal

## Abstract

Processing of negative affective pictures typically leads to desynchronization of alpha-to-beta frequencies (ERD) and synchronization of gamma frequencies (ERS). Given that in predictive coding higher frequencies have been associated with prediction errors, while lower frequencies have been linked to expectations, we tested the hypothesis that alpha-to-beta ERD and gamma ERS induced by aversive pictures are associated with expectations and prediction errors, respectively. We recorded EEG while volunteers were involved in a probabilistically cued affective picture task using three different negative valences to produce expectations and prediction errors. Our data show that alpha-to-beta band activity was related to the expected valence of the stimulus as predicted by a cue. The absolute mismatch of the expected and actual valence, which denotes an absolute prediction error was related to gamma band activity. This demonstrates that top-down predictions and bottom-up prediction errors are represented in specific spectral patterns associated with affective picture processing.

## Introduction

Cortical dynamics induced by emotional picture processing comprise event-related desynchronization (ERD) in the alpha-to-beta band (Cesarei & Codispoti, 2011; Cui et al., 2013; Ferrari et al., 2015; Lee et al., 2017; Meng et al., 2016; Messerotti Benvenuti et al., 2019; Schneider et al., 2019; Schneider et al., 2018; Schubring & Schupp, 2019, 2020) and event-related synchronization (ERS) in the gamma band (Boucher et al., 2015; Güntekin & Tülay, 2014; Keil et al., 2001; Lee et al., 2017; Martini et al., 2012; Müller et al., 1999; Müller et al., 2000; Oya et al., 2002; Schneider et al., 2018; Yang et al., 2020).

Within the framework of predictive coding, lower frequency oscillatory alpha-to-beta band activity has been linked to top-down predictive signals and higher frequency gamma band activity to bottom-up prediction errors (Arnal & Giraud, 2012; Bastos et al., 2012; Huang & Rao, 2011). Predictive coding of perception assumes that neuronal circuits implement perception and learning by constantly matching incoming sensory data with the top-down predictions of an internal or generative model (Clark, 2013; Huang & Rao, 2011; Knill & Pouget, 2004). Consequently, a system can refine models with better predictions by minimizing prediction errors regarding the sensory environment, leading to a more efficient encoding of information (Friston, 2010).

Alpha ERD following affective images is smaller when the image is anticipated, and the tendency is more prominent for negative images (Onoda et al., 2007). On the one hand, these findings suggest that providing a cue prior to an affective image reduces the activation of visual cortex for the image. On the other hand, this might be interpreted as differences in the encoding of expectation signals in a predictive coding framework, where expectation signals manifest as increases in low frequency (alpha-to-beta) activity.

With respect to gamma activity, it has been shown that famous faces elicit a larger gamma response as compared to unfamiliar faces (Anaki et al., 2007; Zion-Golumbic et al., 2010). People see hundreds of unfamiliar faces in daily life, while seeing famous faces is very rare and surprising and leads to a large prediction error. The Free Energy principle including aspects of predictive coding specifically posits the minimization of “free energy” (and thus, indirectly prediction errors) as a mechanism to ensure that agents spend most of their time in a small number of valuable (i.e. positive) states (Friston, 2010). With regards to affective stimuli, this agrees with findings showing that visual stimuli with a negative valence produce larger gamma responses than neutral and positive visual stimuli (Balconi & Lucchiari, 2008; Jung et al., 2011; Keil et al., 2001, 2007; Luo et al., 2007; Martini et al., 2012; Matsumoto et al., 2006; Oya et al., 2002; Sato et al., 2011; Yoshino et al., 2012). Results interpreting the effects of negative valence in the gamma band could be associated with the surprise (i.e. general low probability of a negative encounter) that negative stimuli entail. However, in most studies this cannot be disentangled from the valence as the prediction error associated with a negative stimulus per se cannot be disentangled from the prediction error in the individual experimental setting. To achieve this, additional prediction errors have to be introduced experimentally.

These processes might be interpreted as deficits in learning of the causal structure of events. In predictive coding, an improved causal model by learning improves top-down predictions which consequently lead to a reduction of prediction error signals (Friston, 2012). In this formulation, an important function of emotional valence turns out to regulate the learning rate of the causes of sensory inputs. Specifically it has been proposed that a violation of expectation leads to a (qualitatively) negative valence and increases in the learning rate, while fulfilled expectations are associated with positive valence and a decrease of the learning rate (Joffily & Coricelli, 2013). Absolute prediction errors are also integral part of some learning models. In the Pearce Hall Model (Pearce & Hall, 1980), the absolute error promotes changes in associative strength (i.e. learning rate) such that large absolute prediction errors (surprises) prompt the model to rapidly adapt by increasing its learning rate. If emotions can be derived from a predictive coding function, visual processing of affective pictures can be seen as a simplified model of predictive coding processes in emotion.

In summary, we hypothesize that alpha-to-beta ERD and gamma ERS typically found in responses to negative affective stimuli are actually signals related to predictive coding. This posits that alpha-to-beta ERD responses should be modulated by expectations, whereas gamma ERS responses should be modulated by prediction errors or surprise.

Consequently, we conducted a cue-stimulus paradigm to unravel predictive coding dynamics in affective picture processing. In this paradigm, we restricted presented stimuli to a negative valence and manipulated the degree of negative valence. We expected the anticipated degree of the aversive content to be related to alpha-to-beta ERD. If surprise is a main driving factor of gamma ERS (as derived from a predictive coding perspective), we expected an increase of gamma ERS when there was a mismatch between the anticipated degree of aversion and the actual aversive quality of the picture. If the negative valence or aversive quality is contributing to the gamma ERS effect, we expected an increase of gamma ERS with higher aversion regardless of the anticipated degree of aversion. Finally, based on hypotheses regarding a negative valence associated with prediction errors (Joffily & Coricelli, 2013), we expected a positive modulation of valence ratings by prediction errors.

## Methods

### Participants

We investigated 35 healthy male participants (mean 26, range: 18–37 years), who were paid as compensation for their participation. Applicants were excluded if one of the following exclusion criteria applied: neurological, psychiatric, dermatological diseases, pain conditions, current medication, or substance abuse. All volunteers gave their informed consent. The study was approved by the Ethics board of the Hamburg Medical Association. Data from six participants had to be excluded from the final EEG data analysis due to technical issues during the EEG recording (i.e.: the data of the excluded participants were contaminated with excessive muscle and/or technical artifacts) leaving a final sample of 29 participants.

### Stimuli and Task

Stimulus properties were chosen to be identical to a previous fMRI study of predictive coding where both expectation and absolute prediction error effects were observed in pain (Fazeli and Büchel, 2018).

Aversive pictures were chosen from the International Affective Picture System (IAPS) (Lang et al., 2008) database at three different levels of valence. The images presented during the EEG experiment had three levels of valence of which the low valence category had valence values of 2.02±0.05 (mean ± standard error), the medium valence category had valence values of 4.06±0.02 (mean ± standard error) and the high valence category had valence values of 5.23±0.01 (mean ± standard error). Thermal stimulation was performed using a 30 × 30 mm^2^ Peltier thermode (CHEPS Pathway, Medoc) at three different intensities: low (42°C), medium (46°C), and high (48°C) at the left radial forearm. The pain part of this data will not be described here, but has been reported in Strube et al., 2021.

Prior to each picture or heat stimulus, a visual cue was presented. The color of the cue (triangle, visual angle of each side: 0.96°) indicated (probabilistically) the modality of the stimulus (orange for picture and blue for heat). A white digit depicted inside of each triangle indicated (probabilistically) the intensity of the subsequent stimulus (1, 2 and 3 for low, medium and high valence). During the whole trial, a centered fixation cross (visual angle: 0.24°) was presented on the screen.

Each trial began with the presentation of the cue for 500ms as an indicator for the modality and intensity of the subsequently presented stimulus. The modality (i.e. pain or picture) was correctly cued in 70% of all trials by the color of the triangle. In 60% of all trials, the stimulus intensity was correctly indicated by the digit within the triangle (see Figure 1b for an overview of all cue contingencies).

**Figure 1.**
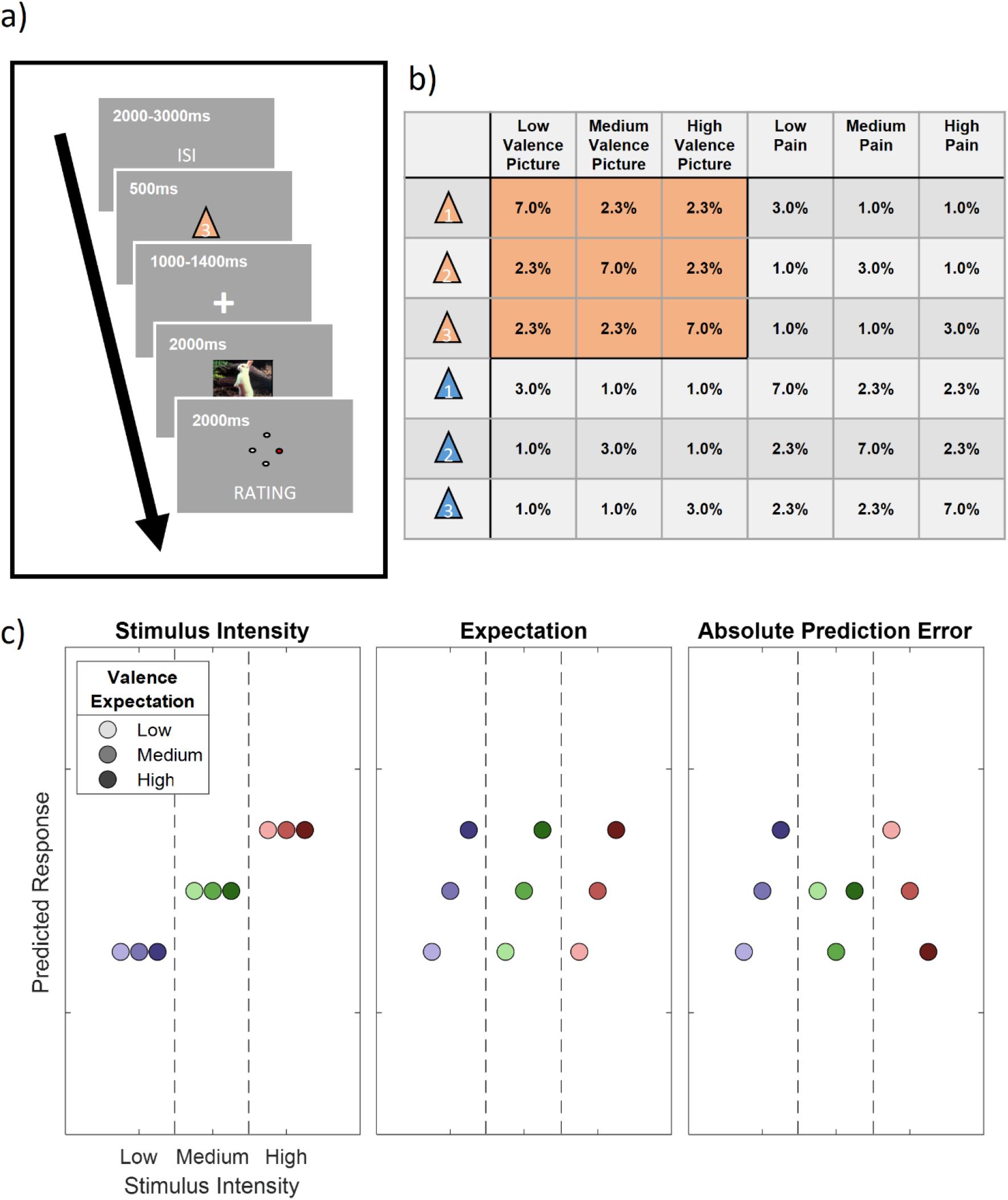
Overview of the study design. a) Graphical representation of the trial structure. Each trial started with the presentation of a cue, indicating the stimulus intensity and modality of the following stimulus. After a jittered phase where only the fixation cross was shown, the stimulus (IAPS picture or pain) was presented. A rating phase (1-4) of the stimulus aversiveness followed. b) Contingency table for all conditions for each cue-stimulus combination. Note that percentages are for all trials, therefore each row adds up to 1/6 (6 different cues). Orange fields indicate conditions included in the analysis, i.e. IAPS pictures where IAPS pictures were indicated by the color of the preceding cue. c) Hypothetical response patterns based on Stimulus Intensity (INT; left), Expectation (EXP; middle) and Absolute Prediction Error (PE; right). The y-axis represents a hypothetical response variable (e.g. EEG power or rating). Each dot represents a different condition for each stimulus-cue combination. Blue colors represent low valence conditions, green colors represent medium valence conditions and red colors represent high valence conditions. Color intensities depict expectation level.

Before the presentation of the stimulus, there was a blank period with a variable duration between 1000ms and 1400ms. The visual (or thermal) stimulus was presented for a duration of two seconds. The visual stimulus (horizontal visual angle of 3.8°; vertical visual angle of 2.4°) was centered on the screen and allowed the participant to perceive it without eye movements. After the termination of the stimulus, subjects were asked to rate the aversiveness of the stimulus on a four point rating scale, where 1 was labeled as “neutral” and 4 was labeled as “very strong”. Ratings were performed using a response box operated with the right hand (see Figure 1a for a visualization of the trial structure).

In addition, four catch trials were included in each block. Subjects were asked to report the preceding cue in terms of their information content of the modality and intensity within 8s and no stimulation was given in these trials.

Trials were presented in four blocks. Each block consisted of 126 trials and four catch trials and lasted about 15 minutes. The trial order within each block was pseudorandomized. The order of blocks was randomized across subjects. The whole EEG experiment including preparation and instructions lasted for about three hours.

Prior to the actual EEG experiment, subjects participated in a behavioral training session. During this session, they were informed about the procedure and gave their written informed consent. The behavioral training session was implemented to avoid learning effects associated with the contingencies between the cues and the stimuli during the EEG session. Between two and three blocks were presented during the training session (without electrophysiological recordings). The experimenter assessed the performance after each block based on the percentage of successful catch trials and the ability to distinguish the three levels of aversiveness of each modality. The training session was terminated after the second block if participants were able to successfully label cues in 75% of the catch trials within the second block.

### EEG Data Acquisition

EEG data were acquired using a 64-channel Ag/AgCl active electrode system (ActiCap64; BrainProducts) placed according to the extended 10–20 system (Klem et al., 1999). Sixty electrodes were used of the most central scalp positions. The EEG was sampled at 500Hz, referenced at FCz and grounded at Iz. For artifact removal, a horizontal, bipolar electrooculogram (EOG) was recorded using two of the remaining electrodes and placing them on the skin approximately 1cm left from the left eye and right from the right eye at the height of the pupils. One vertical electrooculogram was recorded using one of the remaining electrodes centrally approx. 1cm beneath the left eye lid and another electrode was fixated on the neck at the upper part of the left trapezius muscle to record an electromyogram (EMG).

### EEG Preprocessing

The data analysis was performed using the Fieldtrip toolbox for EEG/MEG-analysis (Oostenveld et al., 2011, Donders Institute for Brain, Cognition and Behaviour, Radboud University Nijmegen, the Netherlands. See http://www.ru.nl/neuroimaging/fieldtrip). EEG data were epoched and time-locked to the onset of the IAPS picture. Each epoch was centered (subtraction of the temporal mean) and detrended and included a time range of 3410ms before and 2505ms after trigger onset.

The data were band-pass filtered at 1-100Hz, Butterworth, 4th order. EEG epochs were then visually inspected and trials contaminated by artifacts due to gross movements or technical artifacts were removed. Subsequently, trials contaminated by eye-blinks and movements were corrected using independent component analysis (ICA) (Jung et al., 2000; Makeig et al., 1996). In all datasets, individual eye movements, showing a large EOG channel contribution and a frontal scalp distribution, were clearly seen in the removed independent components. Additionally, time-frequency decomposed ICA data were inspected at a single trial level, after z-transformation (only for artifact detection purposes) based on the mean and the standard deviation across all components separately for each frequency from 31 to 100Hz. Time-Frequency representations were calculated using a sliding window multi-taper analysis with a window of 200ms length, which was shifted over the data with a step size of 20ms with a spectral smoothing of 15 Hz. Artifact components or trials were easily visible and were compared with the raw ICA components. Specifically, single and separate muscle spikes were identified as columns or “clouds” in time-frequency plots. Using this procedure, up to 31 components were removed before remaining non-artefactual components were back-projected and resulted in corrected data. Subsequently, the data was re-referenced to a common average of all EEG channels and the previous reference channel FCz was re-used as a data channel.

Before time-frequency transformations for data analysis were performed on the cleaned data set, the time axis of single trials were shifted to create separate cue-locked and stimulus-locked datasets. Cue-locked data defines the onset of the cue as t = 0. Stimulus-locked data defines t = 0 as the onset of the picture stimulus. Frequencies up to 30 Hz (1 to 30Hz in 1Hz steps) were analyzed using a sliding Hanning-window Fourier transformation with a window length of 300ms and a step-size of 50ms. For the analysis of frequencies higher than 30Hz (31 to 100Hz in 1Hz steps) spectral analyses of the EEG data were performed using a sliding window multi-taper analysis (Strube et al., 2021). A window of 200ms length was shifted over the data with a step size of 50ms with a spectral smoothing of 15 Hz. Spectral estimates were averaged for each subject over trials. Afterwards, a z-baseline correction was performed based on a 500ms baseline before cue onset. For cue-locked data, a time frame ranging from −650ms to −150ms was chosen as a baseline. A distance from the cue onset to the baseline period of 150ms was set because of the half-taper window length of 150ms, i.e. data points between −150ms and 0ms are contaminated by the onset of the cue. For stimulus-locked trials, a variable cue duration (1500-1900ms) was additionally taken into account, resulting in an according baseline from −2550ms to −2050ms from stimulus onset. For the baseline correction of time-frequency data, the mean and standard deviation were estimated for the baseline period (for each subject-channel-frequency combination, separately). The mean spectral estimate of the baseline was then subtracted from each data point, and the resulting baseline-centered values were divided by the baseline standard deviation (Grandchamp & Delorme, 2011).

### Predictive Coding Model

Similar to a previous fMRI study (Fazeli & Büchel, 2018) and our analysis of the pain subset of this dataset (Strube et al., 2021), our full model included three experimental within-subject factors (see Figure 1c). The stimulus intensity factor (INT; see Figure 1c; left column) models the measured response with a simple linear function of the stimulus intensity (−1, 0 and 1 for low, medium and high intensities, respectively). The expectation (EXP) factor was defined (see Figure 1c; center column) linearly from the intensity predicted by the cue. Again, conditions with a low intensity cue were coded with a −1, conditions with a medium intensity cue with a 0 and conditions with a high intensity cue with a 1. The absolute prediction error factor (PE) resulted from the absolute difference of the expectation and actual stimulus intensity (see Figure 1c; right column).

### Behavioral Ratings

Behavioral aversiveness ratings were averaged for all 3×3 cue-stimulus combinations over each participant, resulting in a 29×9 matrix (subject x condition). We tested for main effects across stimulus intensity, expectation, as well as prediction error using a repeated measures ANOVA as implemented in MATLAB (see fitrm and ranova, Matlab version 2020a, The MathWorks).

### EEG data analysis

All statistical tests in electrode space were corrected for multiple comparisons using non-parametrical permutation tests of clusters (Maris & Oostenveld, 2007). Cluster permutation tests take into account that biological processes are not strictly locked to a single frequency or time point and that activity could be picked up by multiple electrodes.

We explored positive and negative time-frequency patterns associated with our variations of stimulus intensity, expectation and absolute prediction errors using a repeated measures ANOVA as implemented in the Fieldtrip toolbox. A statistical value corresponding to p = .05 (F(1,28) = 4.196) obtained from the repeated measures ANOVA for each factor was used for clustering. Samples (exceeding the threshold of F(1,28) = 4.196) were clustered in connected sets on the basis of temporal (i.e. adjacent time points), spatial (i.e. neighboring electrodes) and spectral (i.e. +/− 1Hz) adjacency. Further, clustering was restricted in a way that only samples were included in a cluster which had at least one significant neighbor in electrode space, i.e. at least one neighboring channel also had to exceed the threshold for a sample to be included in the cluster. Neighbors were defined by a template provided by the Fieldtrip toolbox corresponding to the used EEG montage.

Cluster tests were applied separately for low frequencies (1-30Hz in 1 Hz steps) and high frequencies (31-100Hz in 1 Hz steps) in a time frame from 0 (onset of visual stimulus) to 2000ms (end of visual stimulus presentation) for stimulus-locked data and from 0 (onset of cue) to 1500ms (visual stimulus onset) for cue-locked data. Stimulus-locked data was tested for stimulus intensity, expectation and absolute prediction errors factors. Cue-locked data was tested for the expectation factor.

Subsequently, a cluster value was defined as the sum of all statistical values of included samples. Monte Carlo sampling was used to generate 1000 random permutations of the design matrix and statistical tests were repeated in time-frequency space with the random design matrix. The probability of a cluster from the original design matrix (p-value) was calculated by the proportion of random design matrices producing a cluster with a cluster value exceeding the original cluster. This test was applied two-sided for negative and positive clusters.

## Results

### Behavioral data

Participants experienced affective picture (or heat) stimuli which were probabilistically cued in terms of modality and intensity, evoking an expectation of modality and intensity. The subsequently applied stimuli were then rated on a visual analog scale (VAS) from 1-4. Our primary behavioral question was whether ratings are influenced by the experimental manipulation of stimulus intensity, expectation and absolute prediction errors.

To evaluate the main effects of stimulus intensity, expectation and absolute prediction error with regards to the valence of the IAPS pictures, we employed a repeated measures ANOVA of the behavioral data, which revealed significant effects for the main effect of stimulus intensity, i.e. the three levels of valence (F(1,28) = 762.10, p < .001). The main effect for expectation on aversiveness ratings did not yield a significant effect (F(1,28) = 1.46, p = .24). However, the absolute difference between the cued intensity and the actual stimulus intensity (i.e. absolute prediction error), showed a significant effect on aversiveness ratings (F(1,28) = 7.7, p = .01) (Figure 2). In summary, aversiveness ratings were increasing with the degree of aversive valence of the presented picture stimuli. Moreover, these results demonstrate higher ratings when there was a mismatch between the degree of aversion signalized by the preceding cue and the actual stimulus content, i.e. prediction errors are related to higher aversiveness ratings. The results regarding picture and pain stimuli are summarized in Table 1.

**Table 1.** Main effects of stimulus intensity, expectation and absolute prediction errors on subjective aversiveness ratings in picture (visual) and pain (thermal) conditions.

| Factor | Stimulus Intensity |  | Cued Intensity |  | Absolute Prediction Error |  |
| --- | --- | --- | --- | --- | --- | --- |
|  | (INT) |  | (EXP) |  | (PE) |  |
|  | *F*(1,28) | *p* | *F*(1,28) | *p* | *F*(1,28) | *p* |
| **Behavioral Ratings** |  |  |  |  |  |  |
| Thermal | 743.97 | < .001 | 39.53 | < .001 | 2.87 | .10 |
| Visual | 762.10 | < .001 | 1.46 | .24 | 7.7 | .01 |

**Figure 2.**
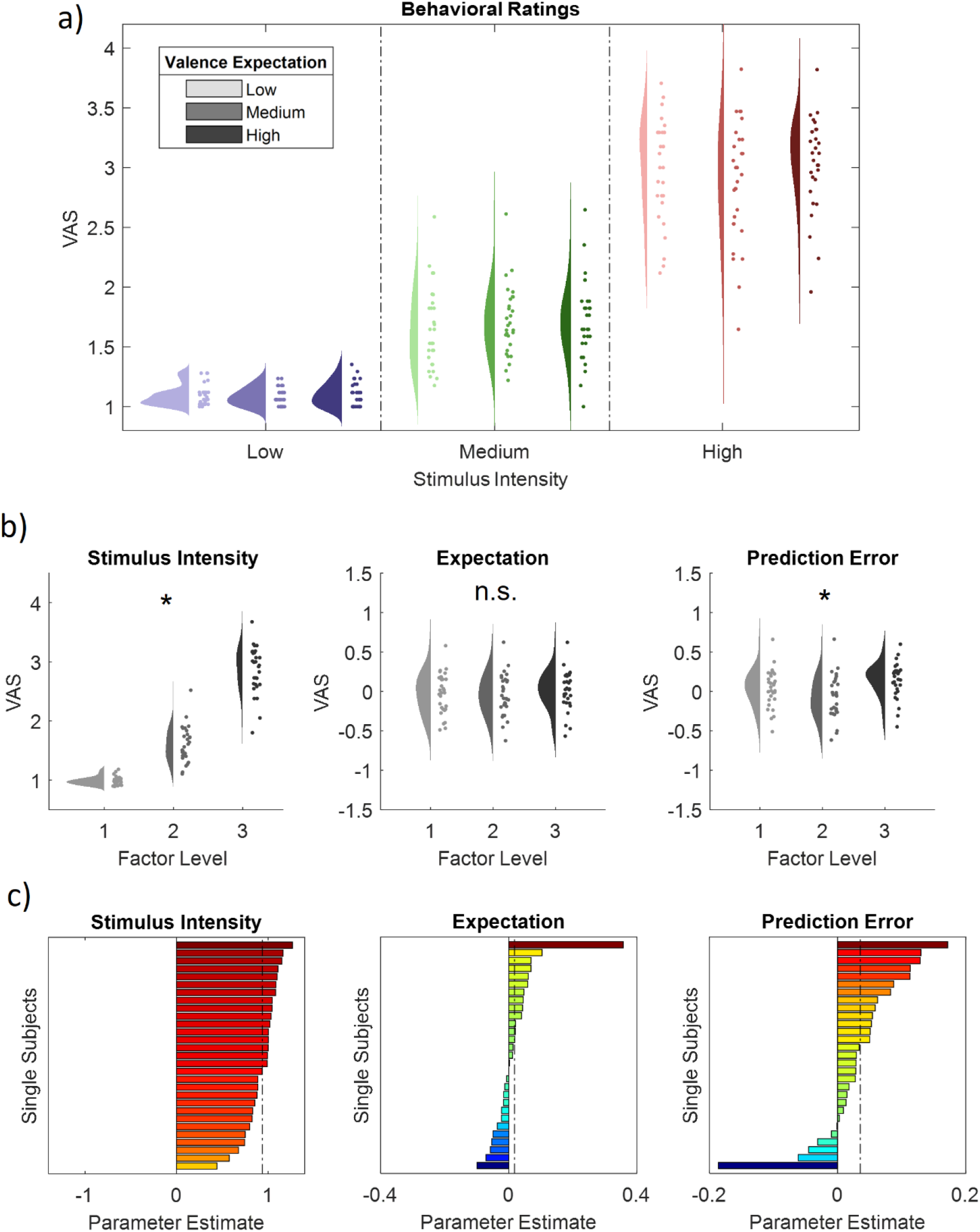
Ratings for IAPS picture stimuli. a) Raincloud plots representing single subject ratings for all 9 congruent conditions (expect a picture and receive a picture). Blue colors represent low valence IAPS picture stimuli, green colors medium valence IAPS picture stimuli and red colors high valence IAPS picture stimuli. Color intensities represent the expected valence from low valence to high valence (low to high color intensities). The data show both an effect of stimulus intensity (increase from blue to green to red), but also a significant positive effect of absolute prediction errors. b) Main effect plots for the stimulus intensity, expectation and prediction error factors (from left to right) showing single subject values and distributions on the response, partialling out (for display purposes) the effects of the other predictors (e.g. EXP and PE were partialled out for the main effect plot of INT) for all three factor levels (increasing from left to right). c) Bars represent the estimated slope for each subject and factor (stimulus intensity, expectation and prediction error from left to right). The dashed line represents the fixed factor estimate (average slope of all subjects). Hot colors represent a positive slope (increases with factor levels) and cold colors a negative slope (decreases with factor levels).

### EEG – Affective Valence

We tested our EEG time-frequency data for a main effect of the valence of the aversive IAPS pictures in the context of a correctly cued modality (i.e. an IAPS picture was expected and received). In order to do so, we performed a repeated measures ANOVA on the time-frequency representation of the EEG data on low frequencies (1-30Hz) and high frequencies (31-100Hz) separately using a cluster correction criterion to address the multiple comparisons problem (see Methods for details). Any significant cluster – composed of neighboring data points in time, frequency and space – would indicate a neuronal oscillatory representation of variations in stimulus intensity in a given frequency band.

In the low frequency (1-30Hz) range, we observed one significant negative cluster of activity (p<.001) indicating a negative linear association of IAPS valence and power in the alpha-to-beta range (Figure 3). Specifically, this negative cluster included samples in a time range from 0 to 2000ms after IAPS stimulus onset in a frequency range from 1 to 30Hz, predominately at frontal and centro-parietal electrode sites. The highest absolute parametric F-value within this cluster obtained from the repeated measures ANOVA F-statistic was F(1,28) = 45.52, (p<.001). This sample was observed at 1150ms after onset of the visual stimulus with a maximum at 21Hz at channel T7. All channels included samples of the negative low frequency stimulus intensity cluster.

**Figure 3.**
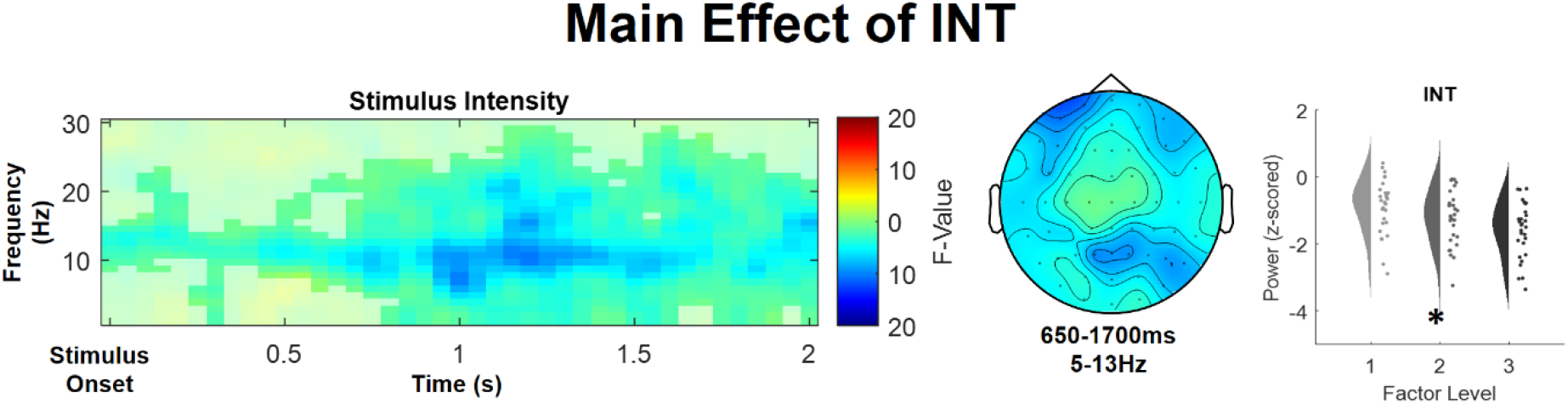
Time-frequency representation (left), topography (center) and main effect plot (right) for the significant stimulus intensity (INT) cluster, showing a decrease of alpha-to-beta power with an increased aversiveness of the stimulus. Time–frequency representations are composed of the statistical F-values of the repeated measures ANOVA averaged over all channels. The significant cluster is highlighted. Hot colors represent a positive slope (increases with factor levels) and cold colors a negative slope (decreases with factor levels). Topographies display the averaged statistical values over time and frequency for each channel. The main effect plot for the INT cluster summarizes single subject values and distributions on the response, partialling out the effects of the respective other predictors (i.e. EXP and PE were averaged out for the main effect plot of INT) for all three factor levels (increasing from left to right).

In conclusion, these results indicate that a higher picture valence is associated with decreased alpha-to-beta band power (ERD) (see Figure 4 for a summary of the results of the main effect of IAPS picture valence). No effect was observed for higher frequencies between 31 and 100Hz.

**Figure 4.**
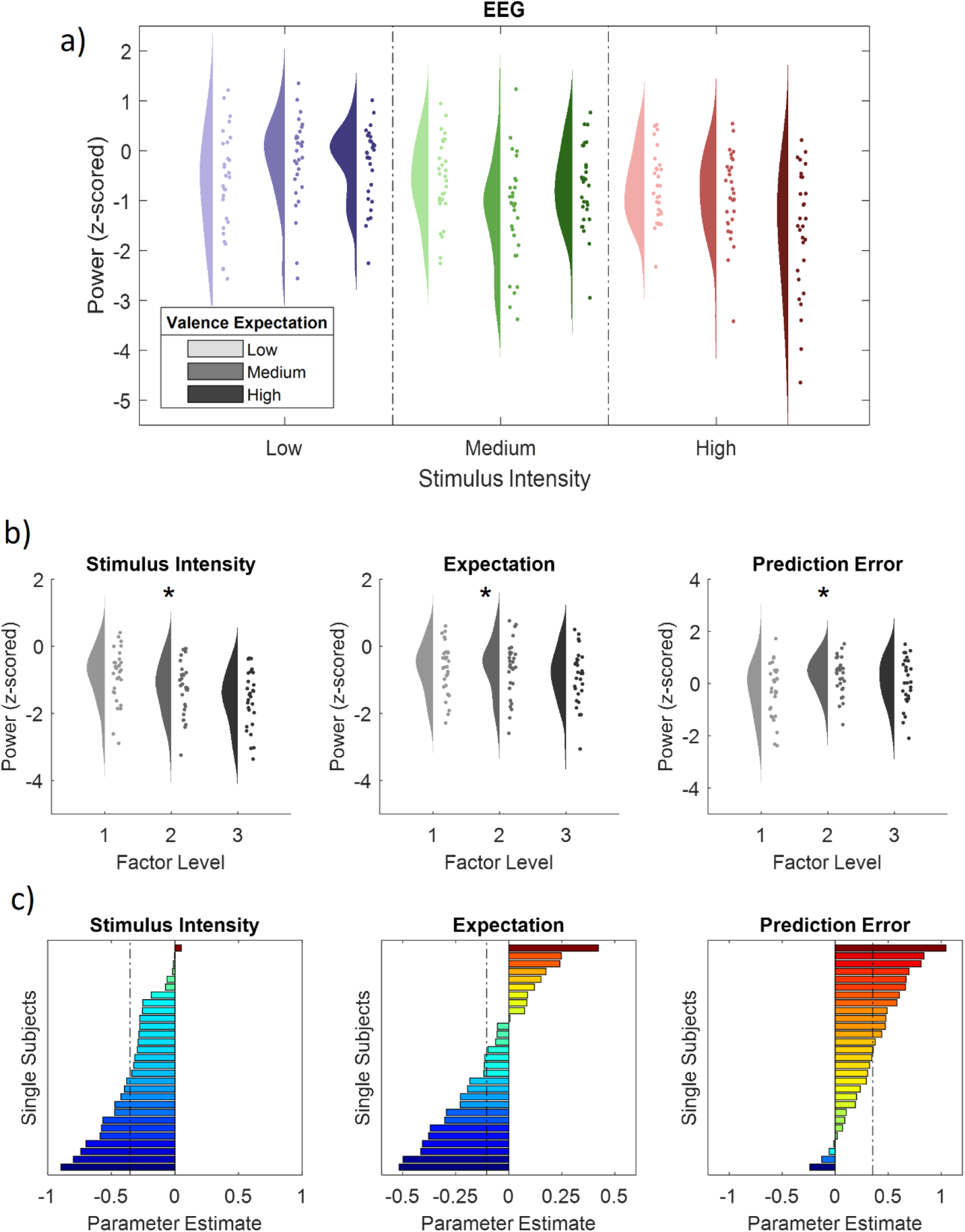
EEG activity at the significant INT cluster related to the manipulation of IAPS valence (INT) and valence expectation (EXP) and the absolute difference of expected and actual stimulus intensities (PE). a) Scatter plots representing single subject EEG power (averaged over all samples included in the significant INT cluster) for all 9 congruent conditions (expect a picture and receive a picture) and according probability distributions averaged over all significant samples included in the negative INT cluster (0-2000ms; 1-30Hz). Blue colors represent low valence IAPS picture stimuli, green colors medium valence IAPS picture stimuli and red colors high valence IAPS picture stimuli. Color intensities represent the expected valence from low valence to high valence (from low to high color intensity). The data shows a significant negative effect of stimulus intensity (decrease from blue to green to red), a significant negative effect of expected valence (decrease from low to high color intensity) but also a significant positive effect of absolute prediction errors. b) Main effect plots for the stimulus intensity, expectation and prediction error factors (from left to right) showing single subject values and distributions on the response, partialling out the effects of the other predictors (e.g. EXP and PE were partialled out for the main effect plot of INT) for all three factor levels (increasing from left to right). c) Bars represent the estimated slope for each subject and factor (stimulus intensity, expectation and prediction error from left to right). The dashed line represents the fixed factor estimate (average slope of all subjects). Hot colors represent a positive slope (increases with factor levels) and cold colors a negative slope (decreases with factor levels).

### Expectation

In a next step, we investigated the representation of the expectation factor (EXP) in our repeated-measures model, again for low frequencies (1–30 Hz) and high frequencies (31–100 Hz) separately in the IAPS stimulus-locked and cue-locked time–frequency representation of the EEG data.

This analysis revealed one significant negative cluster in the low frequency range (1–30 Hz) after IAPS stimulus onset, indicating a negative linear association of cued intensity (EXP) and power in this frequency range (p<0.05). The expectation cluster (p=0.017) included samples from time points ranging from 550 to 1750 ms after IAPS stimulus onset and included frequencies from 3 to 30 Hz. The highest parametric statistical test value (F[1,28] = 30.93, p<0.001) was observed at channel P2 1200 ms after IAPS stimulus onset at a frequency of 20 Hz. All channels included samples of the negative low frequency EXP cluster (see Figure 5).

**Figure 5.**
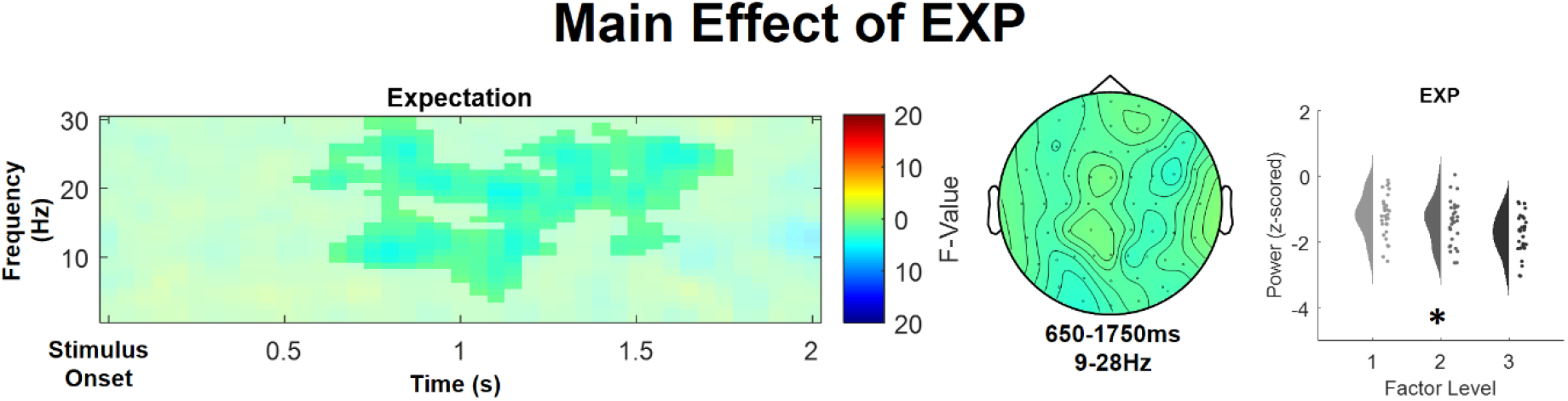
Time-frequency representation (left), topography (center) and main effect plot (right) for the significant EXP cluster, showing a decrease of alpha-to-beta power with an increased anticipated aversiveness of the stimulus. Time–frequency representations are composed of the statistical F-values of the repeated measures ANOVA averaged over all channels. The significant cluster is highlighted. Hot colors represent a positive slope (increases with factor levels) and cold colors a negative slope (decreases with factor levels). Topographies display the averaged statistical values over time and frequency for each channel. The main effect plot for the EXP cluster summarizes single subject values and distributions on the response, partialling out the effects of the respective other predictors (i.e. INT and PE were averaged out for the main effect plot of EXP) for all three factor levels (increasing from left to right).

A cluster analysis of the expectation factor in cue-locked EEG data (from 1 to 30 Hz for low frequencies and 31–100 Hz for gamma frequencies; from 0 to 1500 ms), did not reveal any significant cluster of activity associated with a linear increase or decrease of EXP (all p>0.05).

In conclusion, these results indicate that a higher valence expectation is associated with decreased alpha-to-beta band power during stimulus presentation.

### Absolute Prediction Errors

Finally, we investigated the representation of absolute prediction errors (PE) in our repeated-measures model for low frequencies (1–30 Hz) and high frequencies (31–100 Hz) separately in the IAPS stimulus-locked time–frequency representation of the EEG data. This analysis revealed two significant adjacent positive cluster after IAPS stimulus onset, indicating a linear association of the absolute prediction error factor (PE) and EEG power (p<0.05).

In the high frequency range (31-100Hz) representing gamma activity one positive prediction error cluster was observed (p<.001) and included samples ranging from 0 to 2000ms after IAPS stimulus onset and from 31 to 73Hz. The highest parametric statistical test value (F[1,28] = 61.60, p<0.001) was observed at channel AF7 500ms after IAPS stimulus onset at a frequency of 37 Hz. All channels included samples of the high frequency absolute prediction error cluster (see Figure 6, top row for a summary of the results of the high frequency PE cluster).

**Figure 6.**
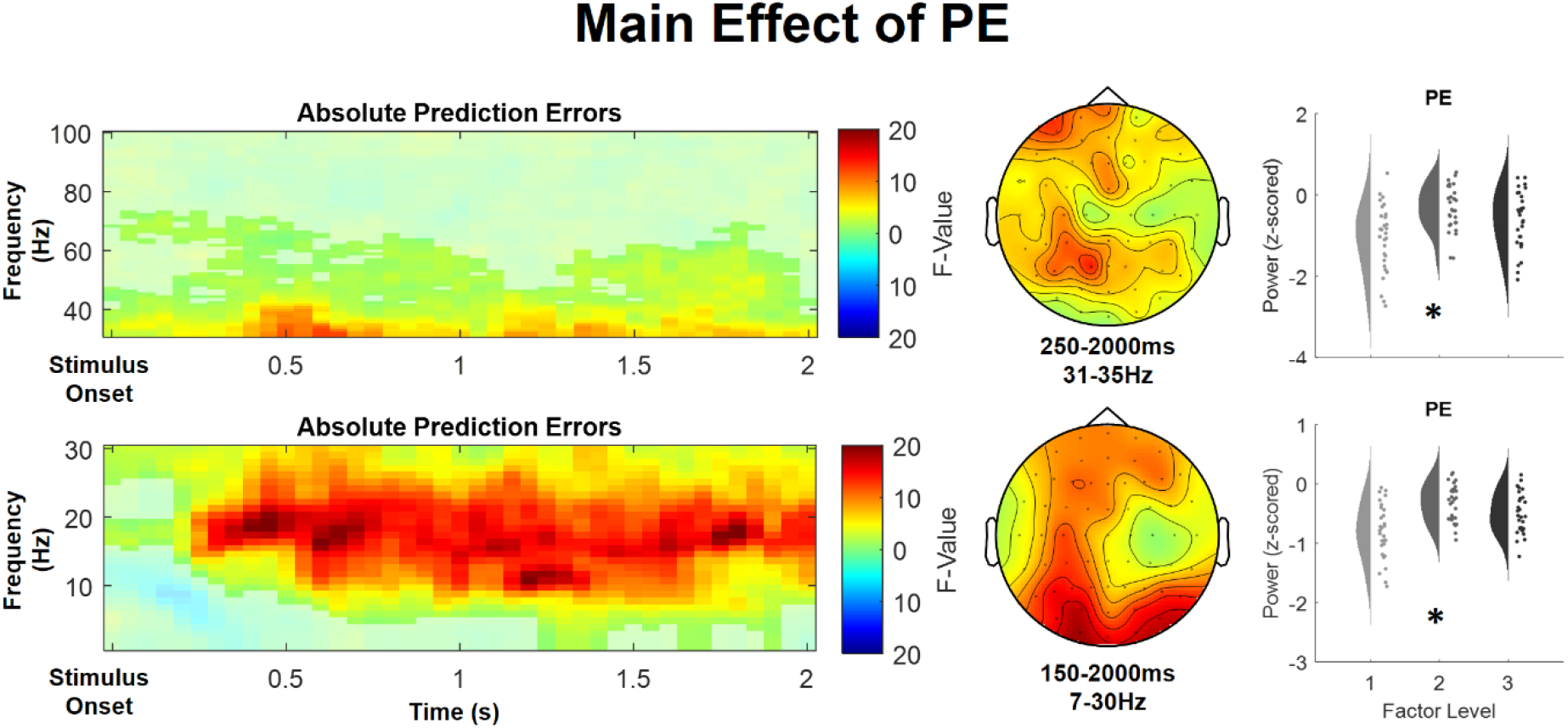
Time-frequency representations (left), topographies (center) and main effect plots (right) for the significant high frequency PE (top) and low frequency PE (bottom) clusters, showing an increase of alpha-to-beta and gamma power with the degree of absolute mismatch of the anticipated and actual aversiveness level of the stimulus. Time–frequency representations are composed of the statistical F-values of the repeated measures ANOVA averaged over all channels. The significant cluster is highlighted. Hot colors represent a positive slope (increases with factor levels) and cold colors a negative slope (decreases with factor levels). Topographies display the averaged statistical values over time and frequency for each channel. Main effect plots for the stimulus intensity, expectation and prediction error clusters summarize single subject values and distributions of the response, partialling out the effects of the respective other predictors (i.e. INT and EXP were averaged out for the respective main effect plot of PE) for all three factor levels (increasing from left to right).

Additionally, one positive prediction error cluster was found in the low frequency range (1-30Hz) (p < 0.001) and included samples from time points ranging from 0 to 2000ms after IAPS stimulus onset and included frequencies from 1 to 30 Hz. The highest parametric statistical test value (F[1,28] = 105.81, p<0.001) was observed at channel PO8 600 ms after IAPS stimulus onset at a frequency of 16 Hz. All channels included samples of the low frequency absolute prediction error cluster (see Figure 6, second row for a summary of the results of the low frequency PE cluster).

In summary, these results suggest an increase in alpha-to-beta and low gamma band power to be associated with expectation violations (i.e. absolute prediction errors), resulting from a mismatch of the cued intensity and the actual valence of the IAPS stimulus.

## Discussion

Using a probabilistic cue paradigm with affective pictures of different valence levels, our data showed a clear discriminability of valence based on behavioral ratings and EEG time frequency patterns. Valence ratings were positively modulated by prediction errors, supporting the hypothesis that prediction errors are linked to higher (negative) valence (Joffily & Coricelli, 2013). With regards to the EEG data, we observed one cluster of activity to be negatively correlated with the valence of the IAPS material in the alpha-to-beta band. Most importantly, our analysis also revealed expectations and violations of expectations (i.e. prediction errors) to be involved in the alpha-to-beta ERD and prediction errors to modulate gamma frequencies.

Alpha-to-beta band activity has been specifically implicated in the processing of top-down expectation signals (Arnal & Giraud, 2012; Bastos et al., 2012). In the processing of affective face stimuli, beta activity has been negatively associated with state anxiety levels in the presentation of affective face stimuli (Schneider et al., 2018). Beta activity has also been linked to top-down prediction signals in the visual perception of causal events (van Pelt et al., 2016).

EEG desynchronization is considered a reliable correlate of excited neural structures or activated cortical areas, while synchronization within the alpha band is hypothesized to be an electrophysiological correlate of deactivated cortical areas (see Pfurtscheller et al., 1996 for a review). An alternative view suggests increased alpha activity to be associated with active inhibition rather than passive inactivity (Foxe & Snyder, 2011; Jensen & Mazaheri, 2010; Klimesch, 2012; Klimesch et al., 2007; Pfurtscheller, 2003; Uusberg et al., 2013). More specifically, it has been suggested that alpha activity represents an attentional suppression mechanism when objects or features need to be specifically ignored or selected against (Foxe & Snyder, 2011). Moreover, event related alpha synchronization is obtained over sites that probably exert top-down control and hence it has been assumed that alpha synchronization reflects a top-down process of inhibitory control (Klimesch et al., 2007).

In this sense, inhibition is a mechanism for gating the flow of information throughout the brain which is mediated by alpha activity (Jensen & Mazaheri, 2010; Klimesch, 2012; Uusberg et al., 2013). In our study, two effects come to play in the alpha-to-beta band, which are relevant with regard to this hypothesis: Firstly, alpha band activity shows a negative relationship with expected stimulus intensity, suggesting less inhibition (i.e. more attention to this information) of highly aversive (potentially negative of threatening) visual stimulation. Secondly, prediction errors resulted in a synchronization of the alpha band, i.e. a positive relationship. In this sense, incongruent trials would be attentionally suppressed and the features would be specifically ignored or selected against. This is because our alpha-to-beta prediction error follows a pattern of higher synchronization with higher prediction errors.

In predictive coding, gamma activity has been specifically associated with prediction error responses (Arnal & Giraud, 2012; Bastos et al., 2012) and has been associated with bottom-up prediction errors in the visual processing of causal events (van Pelt et al., 2016). In the processing of affective stimuli, functional connectivity analyses suggest interhemispheric communication between temporal and frontal regions by phase synchronization at about 40 Hz after stimulus onset (Martini et al., 2012). Intracranial amygdala recordings of patients with chronically implanted depth electrodes revealed that highly arousing and negatively valenced stimuli evoke event-related gamma responses in the amygdala, most pronounced in gamma frequencies between 36-60Hz after stimulus onset (Oya et al., 2002). Moreover, gamma band activity has been found to be positively correlated with state anxiety levels in the presentation of affective face stimuli (Schneider et al., 2018). Our results are in line with these findings, showing an early low-gamma band effect associated with affective picture processing.

In the context of the affective nature of our task, it is interesting to note that participants with dysphoria elicit a smaller alpha ERD in response to pleasant pictures, but not to unpleasant pictures, in frontal-to-parietal electrode sites (Messerotti Benvenuti et al., 2019). In the context of predictive coding, affective disorders (such as major depression) have been linked with bottom-up deficits in predictive processing and increased precision of negative prior beliefs (Kube et al., 2019). The depressed phenotype may emerge from a collection of depressive beliefs associated with the causal structure of the world (Chekroud, 2015). As a consequence, treating depression could be conceptualized as equipping the brain with the resources to modify its internal model of the world (Barrett et al., 2016). Hence, treatment of depression would be associated with brain’s relevant statistical structures becoming less pessimistic (Chekroud, 2015). Thus, the predictive coding model of emotional states associated with affective disorders might be of particular interest for mechanistic insights in depression, as it represents such an internal model with potentially (pathologically) variations in its statistical structure (Barrett et al., 2016; Chekroud, 2015; J. E. Clark et al., 2018; Kube et al., 2020; Smith et al., 2020).

### Limitations

This study has been designed in close analogy to a previous fMRI study to unravel the temporal dynamics of expectation and prediction errors and we decided to use the same experimental paradigm (Fazeli and Büchel, 2018). We therefore decided to also keep the sample characteristics similar and restricted the sample to male participants, which means that we cannot generalize our results to the population. Future studies should investigate samples including female participants. This would also allow to investigate sex effects with respect to expectation and prediction error effects in affective picture processing.

### Summary

Our data show that key variables required for affective picture processing in the context of a generative model (i.e. predictive coding) are correlated with event-related fronto-temporal and centro-parietal alpha-to-beta and gamma activity. These insights shed new light on the role of bottom-up and top-down mechanisms of affective picture processing and suggest predictive coding mechanisms as a possible mechanism to play a role in the processing of affective visual stimuli.

## Acknowledgements

We would also like to thank Matthias Kerkemeyer for his help during data collection. CB is supported by DFG SFB 289 project A02 and ERC-AdG-883892-PainPersist. MR is supported by DFG SFB 289 project A03 and DFG SFB TR 169 project B3. Funded by the Deutsche Forschungsgemeinschaft (DFG, German Research Foundation) – Project-ID 422744262–TRR 289.

